# CD8+ T cells drive myofibroblast activation and contraction via JAK/STAT3 and TGFβ signaling

**DOI:** 10.1101/2025.03.05.641635

**Authors:** Theodoros Ioannis Papadimitriou, Anne van Essen, Merel Willemse, Daphne Dorst, Elly Vitters, Birgitte Walgreen, Hans J.P.M Koenen, Peter van der Kraan, Marije Koenders, Rogier M. Thurlings, Arjan van Caam

**Affiliations:** Department of Rheumatology, Radboudumc, Nijmegen, the Netherlands; Department of Laboratory Medicine-Medical Immunology, Radboudumc, Nijmegen, the Netherlands

**Keywords:** myofibroblast, CD8+ T cells, contraction, fibrosis, cytokine

## Abstract

**Background:** Fibrosis is a major cause of morbidity and mortality in rheumatic connective tissue diseases. In these conditions, fibrotic tissues are characterized by the infiltration of CD8+ T cells and pro-fibrotic myofibroblasts. However, the role of CD8+ T cells in driving myofibroblast activity remains unexplored. Our aim was to investigate this interaction between CD8+ T cells and skin myofibroblasts and to elucidate its underlying mechanisms.

**Methods:** Primary skin-derived myofibroblasts were co-cultured with peripheral blood mononuclear cells (PBMCs) or sorted T cells in a 3D collagen type I hydrogel. To model autoimmunity, allogeneic mismatching was applied. T cell activation and cytokine expression were assessed using flow cytometry and Luminex. Myofibroblast activation was analysed through immunohistochemistry (IHC) and quantitative polymerase chain reaction (qPCR), while activation-induced contraction was measured macroscopically. Intracellular signalling pathway activation in myofibroblasts was evaluated using luciferase reporter cell lines.

**Results:** Co-culture of myofibroblasts with PBMCs strongly induced hydrogel contraction and expression of activation-related markers such as podoplanin, fibroblast activation protein, pSTAT3 and IL-6 in myofibroblasts. CD4+ T cells and CD8+ T cells were also activated by co-culture as identified by increased CD25 and CD69 expression and elevated IL-2 and IFN-γ production. Upon co-culture with either sorted CD4+ or CD8+ T cells, CD8+ T cells more strongly induced myofibroblast contraction and activation than CD4+ T cells. This was not associated with cytotoxicity but with increased IL-6 production by CD8+ T cells compared to CD4+ T cells and STAT3/TGFβ-induced signaling in myofibroblasts. Use of either the JAK/STAT3-inhibitor tofacitinib or the TGFβ receptor inhibitor SB-505124 blocked the activated myofibroblast phenotype, and combined use of both inhibitors had an additive effect on myofibroblast activation and contraction.

**Conclusions:** CD8+ T cells drive primary skin derived myofibroblast contraction and activation not via cytotoxicity-related program but via cytokine release. This sheds light on novel mechanisms of immune cell mediated tissue fibrosis. Furthermore, our results suggest that combining JAK/STAT inhibition with an anti-fibrotic agent that blocks matrix synthesis might be promising in mitigating immune cell mediated tissue fibrosis in connective tissue rheumatic disorders and has added value over blocking these pathways individually.

## Introduction

Fibrosis is characterized by an excessive accumulation of extracellular matrix (ECM) and increased tissue stiffness, leading to loss of tissue architecture and function. Generally, fibrosis is poorly reversible, making it a key complication in various diseases, including (rheumatic) auto-immune diseases such as systemic sclerosis (SSc) (1). Fibrosis has been estimated to be responsible for up to 45% of all deaths in the industrialized world (2). In SSc, fibrosis affects the skin and internal organs such as lungs, gastrointestinal tract and heart leading to a high morbidity and increased mortality (3).

A key cell type in fibrosis is the myofibroblast. This specialized type of fibroblast contributes to fibrosis in multiple ways. For example, myofibroblasts produce large amounts of ECM molecules such as collagen type 1 and fibronectin and make the ECM harder to degrade via expression of collagen crosslinking enzymes and tissue inhibitors of metalloproteinases. Importantly, the defining feature of myofibroblasts is their so-called stress fibers. These fibers, made of alpha smooth muscle actin (*ACTA2*) and (often) non muscle myosin type 2 (NMMII), give the cell contractile properties. Because myofibroblasts are tightly anchored to their environment via cell-cell adherence junctions and cell-matrix focal adhesion junctions, they can contract their environment, leading to tissue stiffness. Notably, increased tissue stiffness precedes matrix deposition in e.g. liver fibrosis (4), indicating that processes increasing tissue stiffness are a crucial initiating step in fibrosis and perpetuating the fibrotic response.

Formation and activation of myofibroblasts can be triggered by the immune system. Cytokines such as TGFβ, IL-4, IL-6 and IL-13 have been described to induce and activate this cell type (5). Various immune cells, such as CD4+ T helper cells, can make these cytokines. Compared to the role of T helper cells in SSc, the role of cytotoxic T cells in myofibroblast formation and activation is less established; typically, these cells are associated with target cell lysis e.g. in anti-viral responses.

We and others recently showed increased numbers of cytotoxic CD8+ T cells in SSc skin and blood compared to healthy controls, especially in early disease processes (6-8). Remarkably, only a small number of CD4+ T helper cells was present in SSc skin, and their presence was similar to that of healthy skin. These observations suggest that CD8+ T cells may exhibit a more pronounced role in connective tissue disease (CTD) fibrosis pathogenesis compared to CD4+ T cells than is currently appreciated. This idea is supported by recent observations of distinct subsets of CD8+ T cells with non-canonical functions divergent from cytotoxicity (9). Such CD8+ T cell subsets have been described to produce T helper cytokines or exhibit regulatory functions (10). Additionally, cytotoxicity-related mechanisms have also been suggested to drive myofibroblast activation via DAMP production resulting from lysis of endothelial cells (11). The question of whether non-canonical CD8+ T cell functions are involved in myofibroblast activation and contraction in CTD, as opposed to cytotoxicity-related processes or CD4+ T helper functions, remains unanswered.

To address this question, we investigated the role of CD4+ versus CD8+ T cells in myofibroblast activation and contraction. For this, we developed a three-dimensional (3D) collagen hydrogel co-culture model comprising primary skin myofibroblasts and alloreactive immune cells. This novel model expands upon classical mixed lymphocyte reaction assays, which have been widely used to evaluate the efficacy and immunogenicity of compounds in vitro and are considered a physiologically relevant approach for studying T cell activation and function. (12). We modified this classical model by incorporating a 3D collagen environment and myofibroblasts to simulate T cell activation, function, and interactions within a fibrotic-like microenvironment, similar to that of the skin. We found that immune cell-mediated myofibroblast contraction and activation was predominantly attributed to cytokine producing CD8+ T cells rather than CD4+ T cells. This process could be halted by inhibiting either JAK/STAT3 or TGFβ signaling. Importantly, combined inhibition showed an additive effect and completely blocked myofibroblast activation and contraction. These data indicate a greater involvement of CD8+ T cells compared to CD4+ T cells in myofibroblast activation, suggesting that therapeutic approaches targeting both CD8+ T cell cytokine production and myofibroblast activation might be beneficial in mitigating immune cell mediated tissue fibrosis.

## Materials and methods

### Isolation and culture of primary skin myofibroblasts

Primary skin dermal myofibroblasts were isolated from 4 mm diameter skin biopsies of the forearm. Written informed consent was provided prior to the procedure (study number: NL57997.091.16). Biopsies were subsequently placed in a 24 wells plate with 2 ml DMEM Glutamax medium (Gibco, Waltham, MA, USA) supplemented with 100 U/ml penicillin, 100 mg/ml streptomycin, 100 mg/L pyruvate, and 20% fetal calf serum (FCS) in standard culture conditions (37 °C, 5% CO-_2_, 95% humidity) for 2 weeks to allow spontaneous outgrowth of primary myofibroblasts. Medium was partly refreshed every 3-4 days. Primary myofibroblasts were cultured on plastic in T175 flasks in DMEM Glutamax medium (Gibco) supplemented with 100 U/ml penicillin, 100 mg/ml streptomycin, 100 mg/L pyruvate, and 10% FCS. Medium was partly refreshed every 3-4 days. Myofibroblasts were used in experiments after passage 6. To validate our findings, Human Dermal Myofibroblasts (HDF) purchased from the American Type Culture Collection (PCS-201-012) were included as a reference strain.

### Isolation and culture of peripheral blood mononuclear cells

Human peripheral blood mononuclear cells (PBMCs) were isolated from buffy coats (obtained from Sanquin, The Netherlands, (project number: NVT 0397-02) by Ficoll Pacque PLUS density centrifugation according to manufacturer’s guidelines. Cells were cultured in RPMI medium 1640 + GlutaMAX™(Gibco) supplemented with 100 U/ml penicillin, 100 mg/ml streptomycin, 100 mg/L pyruvate, and 10% Human Processed Serum (HPS). To aid detection of cytokines with flow cytometry, Brefeldin A (BFA) (5 µg/ml) (Sigma-Aldrich) was added in the culture medium for 3 hours prior to enzymatic digestion (37 °C, 5% CO_2_).

CD3+ T cells, CD4+ T cells and CD8+ T cells were isolated from PBMCs via negative selection with magnetic-activated cell sorting (MACS) kits (Biolegend: 480022, 480010, 480012) according to manufacturer’s guidelines. Following isolation, the enriched fractions of CD3+ T cells, CD4+ T cells, and CD8+ T cells displayed purity levels exceeding 95%, assessed through flow cytometry staining targeting CD3, CD4, and CD8 antigens.

### Myofibroblast and immune cell 3D co-culture hydrogel contraction model

To obtain a single cell suspension, myofibroblasts were washed twice with saline and detached from the culture flask with trypsin/EDTA at 37 °C. After detachment, trypsin was inactivated by adding 10 ml DMEM medium with 10% FCS. Cryopreserved PBMCs were thawed and washed twice with RPMI medium containing 10% HPS.

Next, the cells were seeded into the collagen hydrogel. Every hydrogel consisted of 20 µl Minimum Essential Medium Eagle (Sigma-Aldrich), 10 µl Sodium Bicarbonate 7.5% solution (Gibco), 150 µl Type 1 Bovine Collagen Solution (Advanced BioMatrix) (3.1 mg/ml), and 90 µl cell suspension. The cells in the hydrogel consisted of either 180 000 myofibroblasts alone, 1*10^6^ PBMCs alone, or 180 000 myofibroblasts with different cell concentrations of PBMCs/ CD3+ T/CD4+ T/CD8+ T cells. After the collagen-cell mixture was adequately homogenized, 250 µl of mixture was pipetted per well in a 48 wells plate. The hydrogels solidified for 2 hours in the incubator at 37 °C and 5% CO_2_. After solidification, 750 µl RPMI medium 1640 + GlutaMAX™ (Gibco) supplemented with 100 U/ml penicillin, 100 mg/ml streptomycin, 100 mg/L pyruvate, and 10% HPS was gently pipetted at the side of the wells. For all conditions in every experiment at least three technical replicates were used. Contraction was macroscopically evaluated by scanning the plates on a standard office flat-bed scanner with 600 dpi resolution. The size of the hydrogels was measured using Fiji software and was calculated as follows

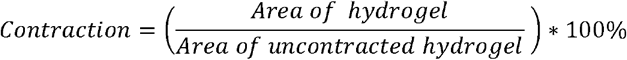

### Pharmaceutical compounds

Tofacitinib (1 µM) (LC Laboratories), SB-505124 (5 µM) (Sigma-Aldrich) were added to the hydrogels by adding the compounds to the medium that was added after solidification of the hydrogels. In the experiments where we wanted to block cytokine production of immune cells, BFA (5 µg/ml) (Merck) was added to isolated immune cells (PBMCs, CD3+ T cells, CD4+ T cells or CD8+ T cells) for 4 hours before being washed away and cells were used for hydrogel co-cultures.

### Enzymatic digestion of hydrogels

To obtain a single cell suspension of the myofibroblasts and PBMCs from the hydrogels in order to study phenotype and gene expression levels of cell populations of interest, hydrogels were enzymatically digested for 1 hour on a roller at 37 °C in 500 µl digestion mix containing collagenase D (Roche) (200 U/ml) and DNase (Roche) (0.1 mg/ml) that were dissolved in PBS. Digestion enzymes were inactivated by adding 50 µl FCS. Hydrogels were then mechanically disrupted by careful pipetting up and down and subsequently spun down for 5 minutes at 300 × g. The supernatant was removed and the cells used for further processing.

### Immunohistochemistry

Collagen hydrogels were formalin-fixed, paraffin-embedded (FFPE), and sectioned into 5.0 µm thick slices before being stained with hematoxylin and eosin (HE). For immunohistochemistry, the FFPE tissue sections underwent deparaffinization using xylene and rehydration with ethanol. Antigen retrieval was performed either in 10 mM sodium citrate buffer (pH 6.0) at room temperature or by heating the slides at 97°C for 10 minutes. Endogenous peroxidase activity was blocked with 3% H-_2_O-_2_ in PBS. Primary antibodies (detailed in **Supplementary Table 1**) and corresponding secondary antibodies (BrightVision Poly-HRP, Immunologic DPVO55HRP, or Envision Flex HRP, DAKO) were applied, followed by labeling with 3,3’-diaminobenzidine (bright DAB, Immunologic or DAKO). The sections were then counterstained with hematoxylin.

### Antibody staining and flow cytometry

Approximately 1 ×10^6^ cells per condition, were washed twice with PBS in a 96 well v-bottom and incubated with (1.5: 1000 PBS) Viakrome808 (Beckman Coulter) viability dye for 30 minutes at 4 °C in the dark. After washing, the extracellular staining was performed by adding 30 µl staining buffer (PBS+1% BSA) buffer with extracellular antibodies for 20 minutes at room temperature protected from light. In case of intracellular staining, cells were fixed with 50 µl Cyto-Fast Fix/Perm solution (Biolegend) for 20 minutes at room temperature. Subsequently, cells were washed twice with 10X Cyto-Fast Perm/Wash Buffer (Biolegend) before stained with intracellular antibodies for 20 minutes at room temperature in the dark. Eventually cells were washed twice and acquired in a Beckman Coulter Cytoflex LX 21-color flow cytometer. Flow cytometry data were analyzed using Kaluza software version 2.1.3 (Beckman Coulter). A list of all extracellular and intracellular antibodies used is provided (**Supplementary table 2, 3**).

For fluorescently activated cell sorting (FACs), cells from digested hydrogels were first filtered through a 70 µm cell strainer (Falcon) and the extracellular staining was performed as previously described while the viability dye efluor780 (eBioscience) was used (1:1000 in PBS). An overview of the gating strategy that was used to sort immune and stromal cell populations of interest is displayed in (**Supplementary figure 1**). Sorted cells were collected in 200 µl RPMI medium supplemented with 10% HPS, 100 mg/L pyruvate, and 1% Penicillin/Streptomycin. After centrifugation, the cell pellet was resuspended in 350 µl RLT buffer from the RNeasy Mini Kit (Qiagen) with 2-Mercaptoethanol (1:100) and stored at -20 °C until further processing.

### RNA isolation and quantitative real-time PCR

RNA isolation was performed with the RNeasy Mini Kit (Qiagen) according to the manufacturer’s manual. RNA concentrations were measured with a nanodrop photo-spectrometer. Amplification grade DNase 1 (Sigma-Aldrich) was incubated for 15 minutes on room temperature in 10X reaction buffer (Sigma-Aldrich). The reaction stopped with Stop Solution For DNase 1 Kit (Sigma-Aldrich) for 15 minutes and by incubation of the samples at 65 °C for 10 minutes subsequently. cDNA was produced with a single step RT-qPCR by the following procedure: Samples were incubated for 5 minutes at 25 °C, for 60 minutes at 39 °C, and for 5 minutes at 95 °C subsequently in a thermocycler. Produced cDNA was diluted 10x with water. To perform qPCR, samples with 5 µl SYBR Green Master Mix (Applied biosystems), 2 µl 1 µM forward and reverse primer solution, and 3 µl cDNA sample were incubated for 10 minutes at 95 °C, then 40 cycli followed including 15 seconds at 95 °C and 60 seconds at 60 °C. After the reaction a melt curve was included in the protocol to verify target specificity. Reference genes used were: *GAPDH, RPS27a*, and *TBP*. Sequences of forward and reverse primers are provided (**Supplementary table 4**). All primers were first validated for their efficacy. -ΔC-_T_ values were calculated with the following formula:

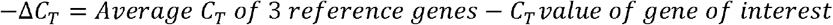

### Reporter Luciferase Assay

Sis-inducible element (SIE) activity was evaluated in reporter constructs cloned in the same primary dermal myofibroblasts and were obtained from Neefjes et al, 2021 (13). Reporter myofibroblasts were also used at a density of 200.000 cells per hydrogel. Recombinant IL-6 (Peprotech: 100-21C-2UG) (10 ng/ml) was used as positive control. After 24 h coculture, hydrogels were collected and centrifuged, after which 150 µl Assay Buffer with 50x substrate from the Nano-Glo Luciferase Assay kit (Promega) was added to the pellet. Luciferase was measured with a Clariostar (BMG Labtech). Emission was measured at 590 nm.

### Apoptosis assay

To differentiate between early apoptotic cells, non-apoptotic cells or cells in late apoptosis/necrosis, single-cell suspensions from digested hydrogels were initially labeled extracellularly with specific monoclonal antibodies for 20 minutes at room temperature. Subsequently, the cells underwent two cold PBS washes and were suspended in 100 µl PBS containing 5 µl 7-AAD (eBioscience, cat# 00-6993-50), 5 µl AnnexinV:FITC labeled (BD Pharmigen), and 0.15 µl CaCl2 (1 M) in PBS. Following a 10-minute incubation in darkness at room temperature, the samples were promptly analyzed via flow cytometry (Cytoflex LX 13, BD) after staining. Cells positive for both 7-AAD and AnnexinV are designated as late apoptotic/necrotic, whereas those positive only for AnnexinV and negative for 7-AAD are considered early apoptotic. Live cells exhibit negativity for both 7-AAD and AnnexinV.

### Cytokine measurements

Human cytokines present in culture supernatants were quantified using Luminex. The Bio-Plex Pro Human Cytokine 27-plex Assay (Bio-Rad, cat# M500KCAF0Y) was employed according to the manufacturer’s guidelines. Analysis of the samples was conducted utilizing BioPlex Manager 4 software (Bio-Rad Laboratories, Hercules, CA, USA).

### Statistical analysis

Statistical analyses were performed with GraphPad Prism version 10.0.3. Pairwise comparisons were analyzed with independent student’s t test. For comparison of two groups a two-sided paired T test was utilized and for comparison of two or more groups a one-way ANOVA was performed. P values below 0.05 were considered to be significant. For multiple comparisons, p values were adjusted for multiple testing with Tukey’s or Dunnett’s multiple comparison test.

## Results

### Co-culture of myofibroblasts with immune cells induces myofibroblast activation and spontaneous contraction

To gain insight into how the interaction between immune cells and myofibroblasts may lead to tissue fibrosis in CTD, we set out to establish a humanized in vitro 3D model to study immune cell mediated myofibroblast activation and contraction. In this model, primary human skin myofibroblasts were co-cultured with allogeneic peripheral blood mononuclear cells (PBMCs) in a collagen type 1 hydrogel (schematic representation of the model is illustrated in **Figure 1A**). Alloreactivity was used to mimic autoreactive immunity in SSc because both are characterized by T cell responses against fibroblast antigens. Co-culturing of PBMCs with myofibroblasts at a 5:1 ratio induced spontaneous hydrogel contraction (**Figure 1B, C**), resulting in an average of approximately 85% within 72 hours (**Figure 1D**). In contrast, myofibroblasts cultured alone showed no contraction (**Figure 1B, C**). The amount and speed of contraction depended on the PBMCs: myofibroblasts ratio (**Supplemental Figure 2**). In addition, we tested multiple primary skin myofibroblast strains with similar effects (**Supplemental Figure 3**).

**Figure 1.**
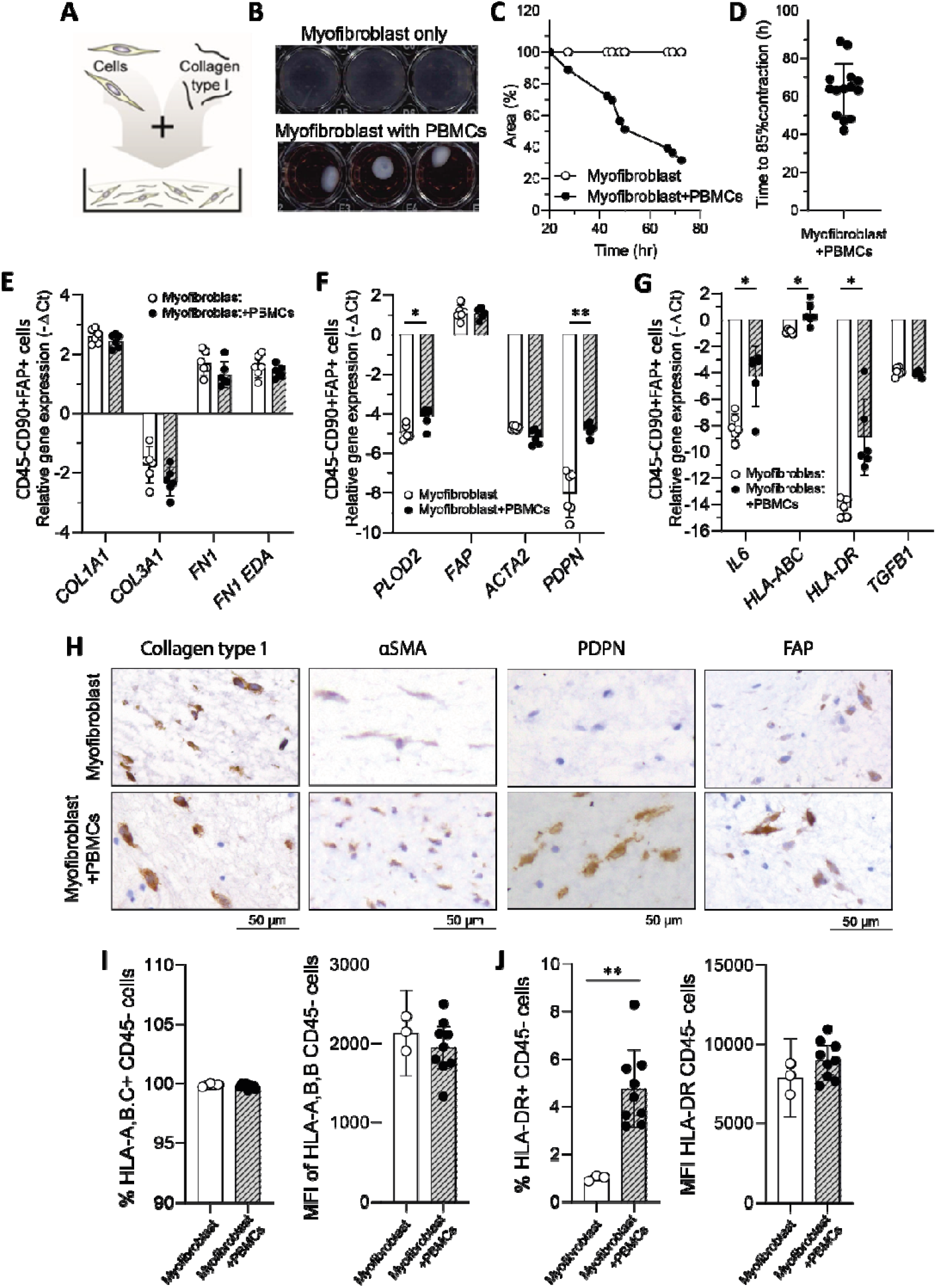
The co-culture of myofibroblasts with immune cells triggers their activation and leads to spontaneous myofibroblast contraction. (A) Schematic illustration of the developed 3D collagen hydrogel contraction model, incorporating co-cultured primary skin myofibroblasts and PBMCs. (B) Representative image exhibiting macroscopic evaluation of myofibroblast contraction in absence or presence of PBMCs. (C) The extent of contraction is quantified relative to the no-cell control and is graphically illustrated for one representative donor over the course of time. (D) Time required until 85% of myofibroblast contraction is reached for each PBMC donor (n=14). (E) Myofibroblasts from collagen hydrogels in absence or presence of PBMCs (n=6 per group) were FACs sorted as the CD45-CD90+FAP+ cell population and analyzed with qPCR for gene expression of markers reflective of (E) ECM production, (F) myofibroblast activation, (G) cytokine production and antigen-presentation capacity. *GAPDH, TBP* and *RPS27A* were used as reference genes. (H) Representative images of Collagen Type I, aSMA, PDPN and FAP immunohistochemistry from collagen hydrogels containing either myofibroblasts alone or myofibroblasts co-cultured with PBMCs. Scale is 50 µm. The percentage of (I) CD45-HLA-A,B,C+ myofibroblasts and HLA-A,B,C mean fluorescence intensity (MFI) (J) CD45-HLA-DR+ myofibroblasts and HLA-DR MFI in the depicted conditions was measured with flow cytometry of the enzymatically digested hydrogels (n=3 myofibroblast only, n=10 myofibroblast + PBMCs).

To better understand immune cell-mediated myofibroblast contraction, we further evaluated the effects of co-culture on myofibroblast biology at both the mRNA and protein levels. For mRNA expression, myofibroblasts were FACS sorted from enzymatically digested hydrogels based on CD45^-^, CD90^+^, and fibroblast activation protein (FAP)^+^ (**Supplemental Figure 1** for the sorting strategy). At the mRNA level, co-culture did not affect the expression of ECM genes such as collagen type I or fibronectin, which are typically upregulated in pathological conditions involving excessive tissue fibrosis, such as SSc. (**Figure 1E**). On the other hand, we observed increased expression of the collagen crosslinking enzyme *PLOD2* (**Figure 1F**), which has been associated with pathological fibrosis (14). Co-culture also strongly enhanced myofibroblast antigen presentation, as it induced the expression of both HLA-ABC (i.e. MHC class 1) and HLA-DRB (i.e. MHC class II) (**Figure 1G**). This suggests that co-culture increases myofibroblast antigen presentation capacity, a pathological process also observed in the context of SSc (15). In addition, co-culture clearly induced *IL6* and podoplanin (*PDPN*) expression, two genes associated with pathologically activated myofibroblasts in e.g. rheumatoid arthritis (**Figure 1G**) (16-18). At the protein level, co-culture notably increased FAP and PDPN expression, as analysed by immunohistochemistry (IHC) (**Figure 1H**). Additionally, the elevated presence of HLA-DR^+^ myofibroblasts was verified at the protein level with flow cytometry (**Figure 1I, J**). Collectively, the developed in vitro model demonstrates that co-culturing skin myofibroblasts with PBMCs induces myofibroblast activation, contraction, and a phenotype with pathological features, highlighting immune-mediated alterations in myofibroblasts.

### Co-culture of immune cells with myofibroblasts induces T cell activation, cytotoxicity and cytokine production

To examine the influence of myofibroblasts on the co-cultured immune cells, we performed multi-color flow cytometry on the single cells isolated after digestion of the collagen hydrogels at multiple timepoints (24, 48, 72 hours). Since our model was based on allogeneic mismatch, we evaluated the effect of the co-culture on both CD4^+^ T cell and CD8^+^ T cell activation. To assess this, we measured T cell activation markers (CD25, CD69, CD134) after 16 hours of co-culture and compared them to immune cells cultured in the collagen hydrogel without myofibroblasts. CD25 and CD69 were significantly induced in both T cell subtypes, while no difference in CD134 (OX40) expression was observed (**Figure 2A**). Since cytokine production is a result of T cell activation, we examined the kinetics of T cell activation over extended culture periods by measuring IL-2 and IFNγ expression for up to three days of co-culture. Initially, approximately 10% of T cells were positive for these cytokines, but the proportion of IL-2 and IFNγ-expressing CD4+ and CD8+ T cells increased over time, reaching up to 80% at 72 hours (**Figure 2B, C**). Additionally, CD8+ T cells showed a time dependent elevation of granzyme B expression that fits with their increased activation status (**Figure 2D**). In conclusion, myofibroblast activation, contraction, and pathogenic phenotype are accompanied by the induction of effector functions in both CD4+ and CD8+ activated T cells. Thus, our model extends beyond myofibroblast activation.

**Figure 2.**
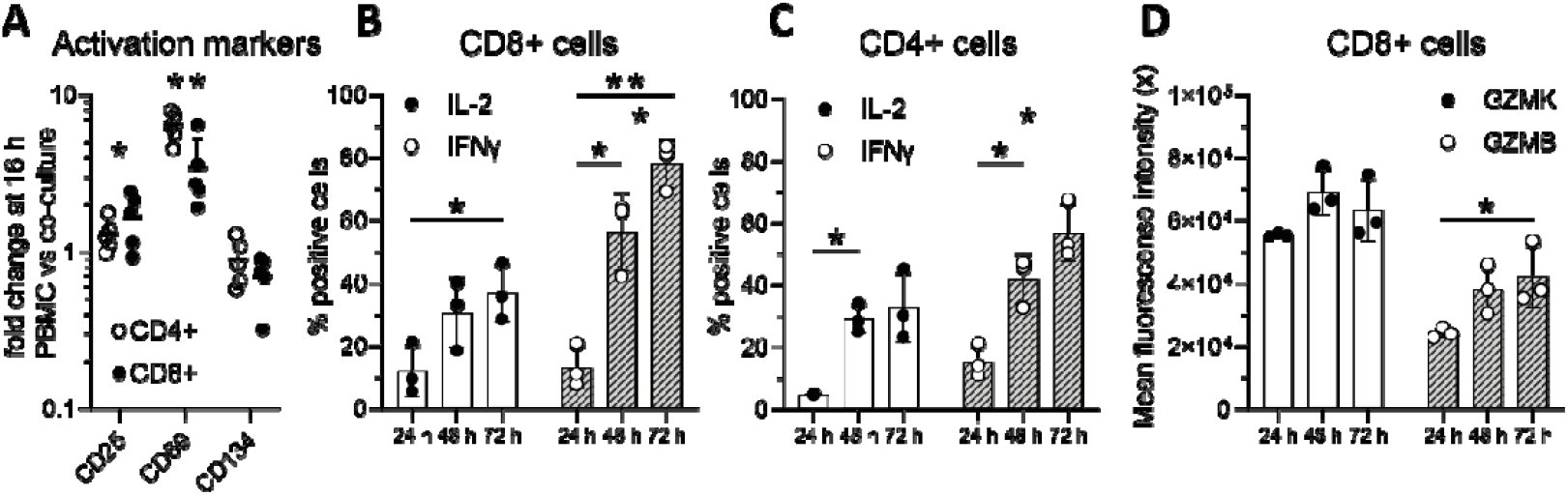
Co-culturing immune cells with myofibroblasts promotes T cell activation, granzyme expression, and cytokine production. (A) The expression of the activation markers CD25, CD69, CD134 on both CD4+ and CD8+ T cells was measured with flow cytometry of enzymatically digested myofibroblast and PBMC (n=5) hydrogels after 16 hours of co-culture. Values are represented as fold change compared to PBMCs cultured alone. Asterisks refer to both CD4+ and CD8+ T cell fold change statistical significance compared to PBMCs alone condition. Flow cytometric quantification of the percentage of IL-2 and IFNγ positive CD8+ T cells (B) and CD4+ T cells (C) after 24, 48 and 72 hours of co-culture (n=3). (D) Mean fluorescence intensity of the expression of granzyme K (GZMK) and granzyme B (GZMB) in CD8+ T cells after 24, 48 and 72 hours of co-culture (n=3).

### Myofibroblast contraction induced by immune cells is not driven by cytotoxic mechanisms

Given that elevated myofibroblast contraction was linked to T cell activation, we next investigated whether this could be attributed to increased T cell cytotoxicity towards myofibroblasts as target cells. To determine if the enhanced contraction was associated with cell death or cell death-related fragments (which can trigger myofibroblast activation), we quantified myofibroblast cell death using flow cytometry and IHC. First, we assessed myofibroblast viability using a fixable viability dye in flow cytometric staining of single cells from the digested hydrogels. The presence of PBMCs did not negatively affect myofibroblast viability (**Figure 3A**). On IHC, a limited number of apoptotic cells expressing active caspase 3 were observed, and only a few cells showed an increased amount of double-stranded DNA breaks (γH2AX) when comparing myofibroblasts cultured alone to those co-cultured with PBMCs (**Figure 3B**). Further investigation using 7AAD and Annexin-V flow cytometric staining to measure early and late apoptosis, as well as necrosis of myofibroblasts, revealed only a small percentage (<5%) of dead or pre-apoptotic myofibroblasts in both short-term (24 hours) and long-term (72 hours) cultures (**Figure 3C-F**). Collectively, our data suggest that immune cell-mediated myofibroblast cell death is minimal and does not appear to be the primary mechanism driving myofibroblast activation and contraction.

**Figure 3.**
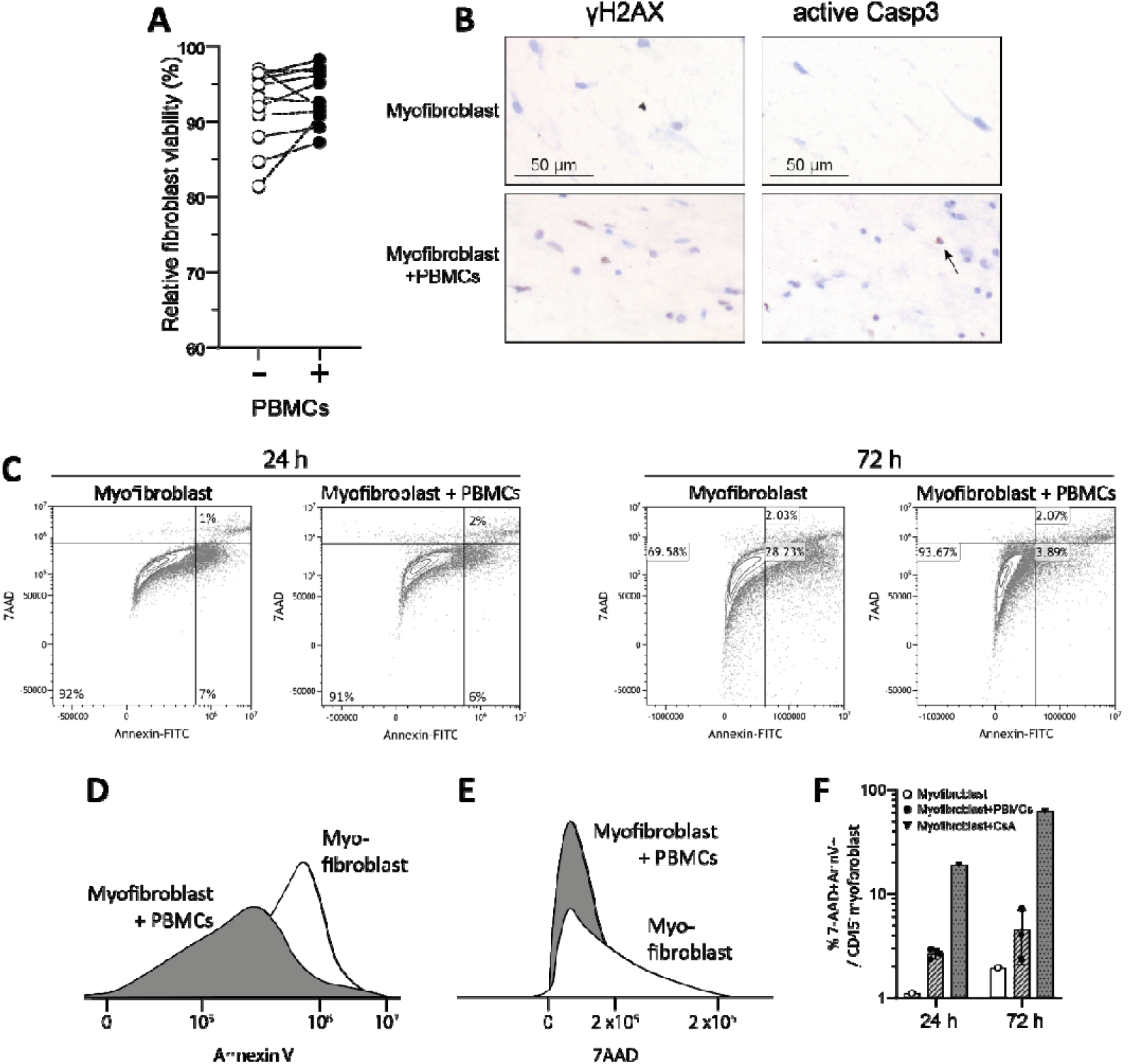
Immune cell-induced myofibroblast contraction is not dependent on cytotoxic mechanisms. (A) Percentage of relative myofibroblast viability in absence or presence of PBMCs, measured by flow cytometric viability staining. (B) Representative images of γH2AX and active caspase-3 immunohistochemistry from collagen hydrogels containing either myofibroblasts alone or myofibroblasts co-cultured with PBMCs. Scale is 50 µm. (C) Representative flow cytometry plots illustrating myofibroblast 7-AAD/ Annexin V staining at 24 and 72 hours of co-culture. Flow cytometry overlay histograms of one representative experiment comparing mean fluorescence intensity of (D) Annexin V and (E) 7-AAD staining between myofibroblast and myofibroblast+ PBMC conditions at 72 hrs. (F) Percentage of 7-AAD+/ AnnexinV+ CD45-myofibroblasts at 24 and 72 hours of cell culture alone or in presence of PBMCs (n=3). A condition with myofibroblasts treated with cyclosporin A (CsA) was used as positive control.

### Cytokine producing CD8+ T cells drive myofibroblast contraction

Recent data suggests a prevalent presence of CD8+ cytotoxic compared to CD4+ T helper cells in the affected tissues (skin, lungs, synovium) of patients with autoimmune mediated CTD (6, 7, 19). Therefore, we analyzed whether CD8+ T cells and CD4+ T cells were differentially linked to the observed myofibroblast contraction. To address this, we sorted CD3+ T cells, CD4+ T cells and CD8+ T cells through negative magnetic beads selection from healthy blood and co-cultured them in the same cell concentration separately with myofibroblasts. All T cell populations induced myofibroblast contraction, and of note, CD8+ T cells induced significantly higher contraction compared to CD4+ T cells (**Figure 4A**). To further evaluate whether TCR signaling is required for the observed contraction, we used an anti-CD8 blocking antibody. Blocking the CD8 receptor resulted in a 25% decrease in myofibroblast contraction but did not completely inhibit it (**Figure 4B**), suggesting that mechanisms downstream of TCR activation are not the only way CD8+ T cells activate myofibroblasts. By transferring the supernatant from the myofibroblast: immune cell co-culture, we were able to trigger hydrogel contraction when added to new, unexposed hydrogel mono-cultured myofibroblasts. This suggests that soluble mediators, such as cytokines, are involved (**Supplemental Figure 4**). The supernatant was screened for a large panel of cytokines, detecting IL-6, IL-8 and IFNγ, while IL-1β, IL-2, IL-4, IL-5, IL-17, IL-12, IL-13, IL-10, IL-15, and IL-17 were undetectable.

**Figure 4.**
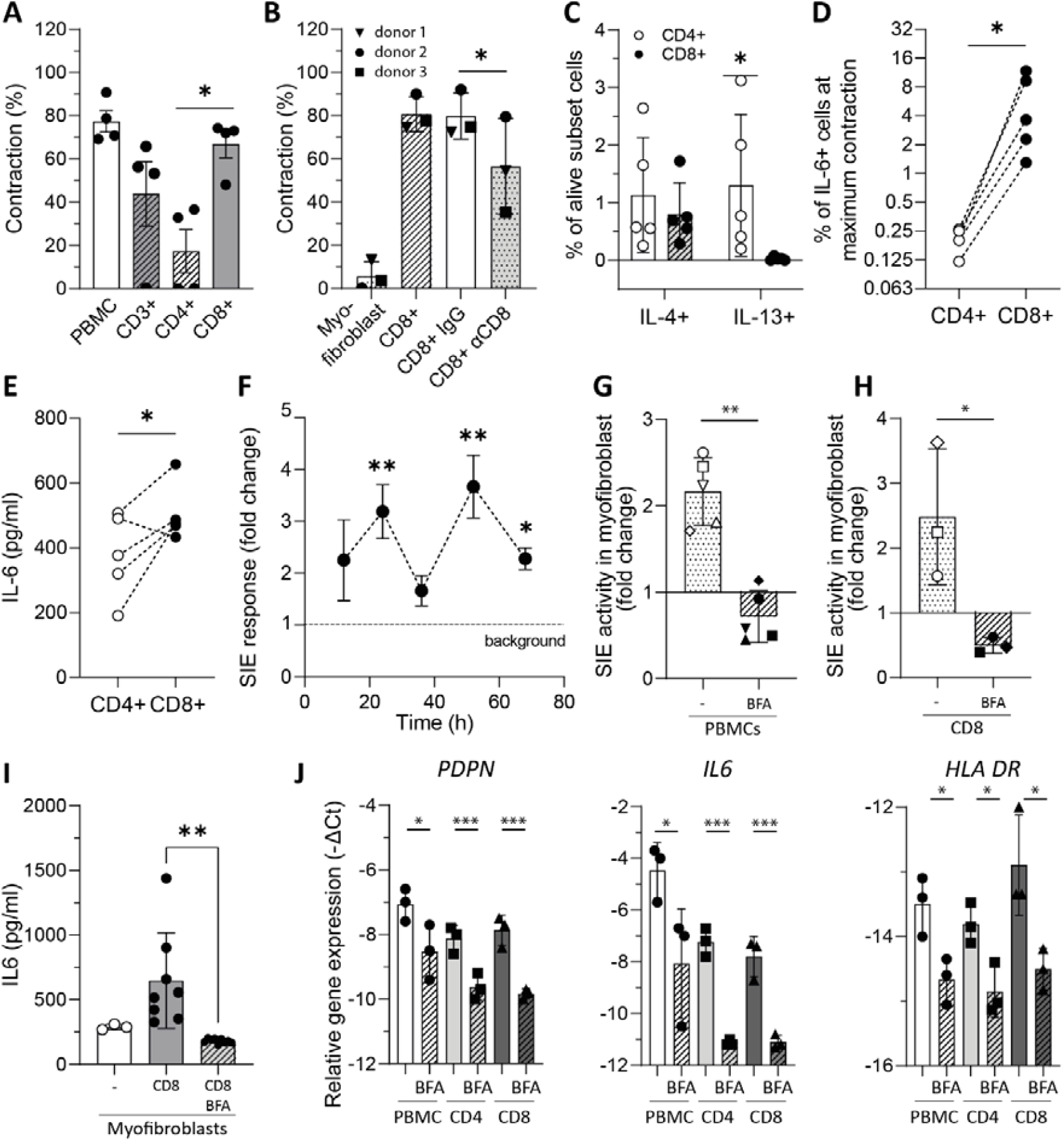
Myofibroblast contraction is driven by cytokine-secreting CD8+ T cells. (A) Level of myofibroblast contraction after co-culture with PBMCs or sorted CD3+/CD4+/CD8+ T cells (n=4) at time of maximum contraction. (B) Level of myofibroblast contraction after co-culture with CD8+ T cells in presence/absence of anti-CD8 monoclonal antibody or isotype control (n=3). Expression of IL-4 and IL-13 (C) and IL-6 (D) on CD4+ and CD8+ T cells was measured with intracellular flow cytometry of digested hydrogels containing PBMCs and myofibroblasts (n=5) at time of maximum contraction. (E) Production of IL-6 in the co-culture supernatant of myofibroblasts that were co-cultured with either CD4+ or CD8+ T cells. (F) SIE response of primary myofibroblast cell line was evaluated with the luciferase reporter assay after cells were incubated with supernatant from hydrogels containing myofibroblasts and PBMCs (n=3) that was collected at the depicted time points. Fold change of SIE activity in primary myofibroblast cell line incubated with supernatant from hydrogels containing myofibroblasts and (G) PBMCs or (H) CD8+ T cells that were co-cultured for 24 hours. In depicted experimental conditions BFA was added to block release of cytokines. (I) Total IL-6 levels of co-cultures after 24 hours. (J) Primary skin myofibroblasts were cultured for 24 hours with cell culture supernatants obtained from myofibroblast hydrogels containing PBMCs, CD4+ T cells, or CD8+ T cells in absence or presence of BFA. Myofibroblast relative gene expression of PDPN, IL-6 and HLA-DR was measured with qPCR and values are represented as -ΔCt. PBMCs; Peripheral Blood Mononuclear Cells, SIE; Sis-Inducible Element, BFA; Brefeldin A, PDPN; Podoplanin

Given that supernatant transfer induced myofibroblast contraction in the absence of immune cells and in light of the recently described T helper cytokine production by CD8+ T cells (10), we compared the expression of T helper cytokines (i.e., IL-4, IL-13, IL-6, and IFNγ) between CD4+ and CD8+ T cells to better understand their potential involvement in our model. We found that IL-4 was produced by both T cell subtypes, while IL-13 was only produced by CD4+ T cells (**Figure 4C**). In contrast to previous research on the pro-fibrotic role of these cytokines in SSc and other CTDs, it seems unlikely that they are the main drivers of the CD8+ T cell effects in this model. However, both the expression (**Figure 4D**) and production (**Figure 4E**) of the pro-inflammatory cytokine IL-6 were significantly elevated in CD8+ compared to CD4+ T cells, suggesting a potential role in the enhanced myofibroblast contraction.

STAT3 is a prominent intracellular mediator of IL-6 signaling. Therefore, we next used a luciferase construct to measure activity of this transcription factor: i.e. sis-induced element (SIE) driven luciferase expression. Co-culture of PBMCs with these reporter cells resulted in clear SIE-driven luciferase expression. Interestingly, we observed an oscillating SIE-driven luciferase expression reflecting a dynamic regulation over time (**Figure 4F**). We then sought to evaluate whether the elevated SIE response was attributed to cytokine production from PBMCs and CD8+ T cells. To do this, we pre-incubated whole PBMCs or isolated CD8+ T cells with the Golgi inhibitor BFA to block their cytokine production and then co-cultured them with myofibroblasts carrying the SIE luciferase construct. Cytokine blockade in both PBMCs and CD8+ T cells fully inhibited SIE activity (**Figure 4G, H**), showing that the elevated SIE-driven myofibroblast activity was likely mediated by a secreted factor, such as IL-6. This hypothesis was confirmed by measuring IL-6 levels in the supernatant of these cultures, where pre-treatment of CD8+ T cells with BFA completely blocked IL-6 production (**Figure 4I**). Furthermore, blocking cytokine production in CD8+ T cell also halted myofibroblast gene expression of PDPN, IL-6 and HLA-DR (**Figure 4J**). In conclusion, the increased myofibroblast activation and contraction observed in our model is a dynamic process that requires interaction between cells and cytokines, and it appears to be predominantly driven by CD8+ T cells in a STAT3 signaling-dependent manner.

### Co-culture mediated myofibroblast activation is dependent on JAK/STAT and TGFβ signaling

To further unravel the molecular mechanisms underlying STAT3 CD8+ (IL-6+) T cell-mediated myofibroblast contraction, we next evaluated myofibroblast expression of key downstream TGFβ and JAK/STAT signaling molecules (pSMAD2 and pSTAT3, respectively) 72 hours after their co-culture with PBMCs. Both SMAD2 and STAT3 phosphorylation were highly activated in the presence of immune cells (**Figure 5A**). Interestingly, the expression of these markers was reciprocal, suggesting distinct myofibroblast mechanisms and functions.

**Figure 5.**
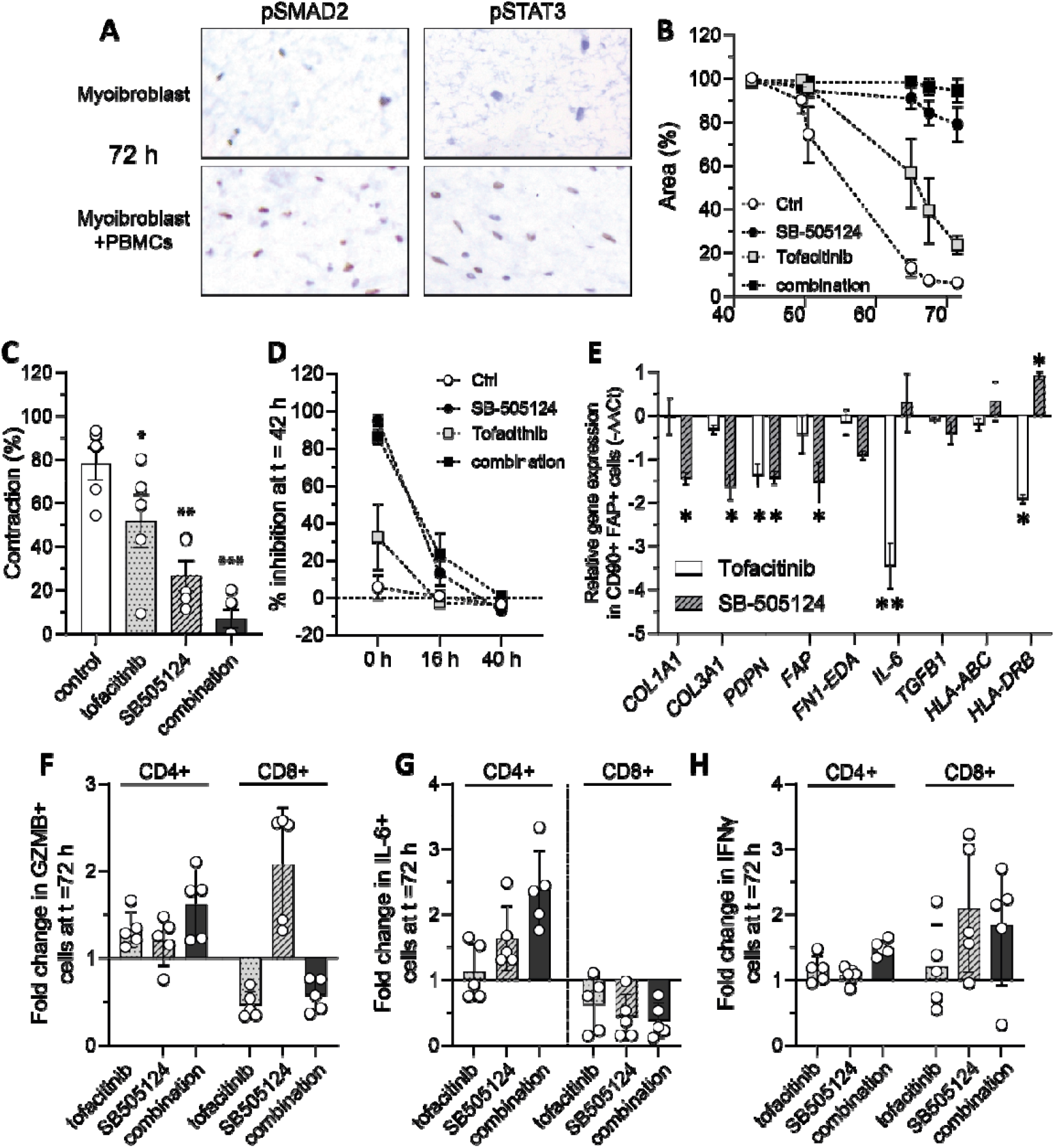
CD8+ T cell mediated myofibroblast activation is dependent on JAK/STAT and TGFβ signaling. (A) Representative images of pSMAD2 and pSTAT3 immunohistochemistry from collagen hydrogels containing either myofibroblasts alone or myofibroblasts co-cultured with PBMCs. Scale is 50 µm. (B) Level of myofibroblast contraction in myofibroblast: PBMCs hydrogels (n=5) that were treated with SB-505124 inhibitor, tofacitinib or a combination of the inhibitors (n=5). Graph represents mean SD myofibroblast contraction over time and the individual values at time of maximum contraction are depicted in panel (C). (D) Percentage of inhibition in myofibroblast contraction after 42 hours treatment with the indicated inhibitors (n=5). Values are mean SD. In this experiment inhibitors were added at the beginning (t=0) or after 16 or 40 hours of culture. (E) Relative gene expression (−ΔΔCt) of selected genes in sorted CD90+FAP+ myofibroblasts from PBMC/myofibroblast hydrogels that were treated with tofacitinib or SB-505124. From the same hydrogels, digested cells were subjected to flow cytometry staining to evaluate the effect of the inhibitors towards expression of (F) GZMB (G) IL-6 and (H) IFNγ in both CD4+ and CD8+ T cells. Results are represented as fold change compared to control condition (n=5). PBMCs; Peripheral Blood Mononuclear Cells,

To assess the potential reversibility of these processes, we treated PBMC: myofibroblast hydrogels with the JAK/STAT inhibitor tofacitinib and the TGF-β1 receptor (ALK4/5/7) inhibitor SB-505124 to evaluate their effects on myofibroblast contraction. Neither inhibitor negatively affected myofibroblast or immune cell viability (**Supplemental Figure 5**), but both significantly reduced myofibroblast contraction. Furthermore, the combination of both inhibitors exhibited an additive effect, almost completely preventing myofibroblast activation (**Figure 5B, C**). These results suggest that TGFβ and JAK/STAT signaling play key roles in mediating the elevated myofibroblast activation.

To further investigate the kinetics of myofibroblast contraction inhibition by these inhibitors, we introduced them into the co-culture either at baseline or 16 and 40 hours after initiation. The addition of the inhibitors at 16 hours still showed an effect; however, when added after 40 hours, no effect was observed (**Figure 5D**). This suggests that the underlying molecular mechanisms occur early in the model and are not reversible with later intervention.

In addition to the (kinetic) effect of the inhibition on myofibroblast contraction, we were also interested in its potential effect towards other myofibroblast functions such as ECM production, antigen presentation and activation. To address this, we analyzed the gene expression of FACs sorted CD45-CD90+FAP+ myofibroblasts that were treated or untreated with the aforementioned inhibitors. SB-505124 effectively halted pro-fibrotic myofibroblast phenotype by decreasing the expression of ECM and activation genes such as *COL1A1, COL3A1, PDPN* and *FAP*, while tofacitinib showed a significant reduction of myofibroblast *IL-6* and *HLA-DRB* expression.

Both inhibitors may also directly or indirectly affect immune cell function. Thus, we finally analyzed the effects of these inhibitors on CD8+ T cells and CD4+ T cells. The inhibitors did not significant affect CD4+ T cell cytokine production (IL-6 and IFNγ) (**Figure 5F-H**). Of note, tofacitinib, but not SB-505124, inhibited CD8+ T cell cytotoxic mediators, as indicated by reduced GZMB expression (**Figure 5F**). These findings further support the notion that CD8+ T cell, but not CD4+ T cell, cytokine production is driving myofibroblast activation. Interestingly, both tofacitinib and SB-505124 significantly reduced IL-6 production, but not IL-4, IL-13, or IFNγ, in CD8+ T cells (**Figure 5G**). In conclusion, these results suggest that myofibroblast activation depends on non-canonical CD8+ T cells producing IL-6, rather than exhibiting direct cytotoxicity, and that this activation can be reversed by targeting distinct immune cell and myofibroblast molecular pathways. Targeting both pathways shows an additive effect in halting T cell effector functions, myofibroblast activation, and contraction.

## Discussion

Collagen contraction assays serve as models for tissue contraction, utilizing the observation that collagen hydrogels populated with cells undergo predictable and consistent contraction over time (20). In this study, we expanded upon current models by introducing allogeneic mismatching to mimic the interaction between auto-reactive immune cells and myofibroblasts, aiming to investigate how adaptive immune cells drive tissue contraction. Strikingly, CD8+ T cells were more effective at inducing myofibroblast contraction than CD4+ T cells, and this effect does not appear to be driven by cytotoxic mechanisms. Further analysis revealed that myofibroblast activation was dependent on JAK/STAT3 and TGFβ signaling pathways, which function differently in CD4+ and CD8+ T cells. Notably, simultaneously targeting these pathways had an additive effect in halting hydrogel contraction, likely by inhibiting both T cell and myofibroblast activation.

We developed a novel 3D hydrogel co-culture model that induces spontaneous myofibroblast contraction through allogeneic immune cells. This model builds upon the classic mixed lymphocyte reaction model and a 3D myofibroblast contractility model. Our findings show that this model recapitulates key features of pathogenic myofibroblasts, including tissue contraction, expression of cross-linking enzymes, and activation and antigen-presentation. Co-culture strongly increased expression of activation markers such as FAP, PDPN, HLA type 2 and IL-6—markers of a myofibroblast phenotype associated with chronic inflammation, as seen in conditions like rheumatoid arthritis, and indicative of immune crosstalk (21). However, no significant changes were observed in ECM molecules. Future studies with extended culture times could further refine the model, providing deeper insights into immune cell-induced myofibroblast ECM production and deposition.

Mixed lymphocyte T cell alloreactivity depends on direct recognition of mismatched MHC class I or II molecules and/or of (allo)peptides presented by these MHC molecules (22). In mixed lymphocyte cultures, as well as, in vivo, approximately 5-10% of the T cell pool becomes activated in a highly polyclonal manner (12, 23) with a similar response for CD4+ and CD8+ T cell populations (24). In our experiments, expression of the classical T cell activation markers (25) CD25 (IL2R) and CD69 was strongly induced in both CD4+ and CD8+ populations after 24 hours of co-culture, but expression of CD134 (OX40) was not. However, kinetics of OX40 expression are slower than that of CD25 and CD69 and typically peak after 48 hours (25). Based on IL-2 and IFNγ expression, we observed activation of 5-10% of CD4+ and CD8+ T cells similar as reported before, but these numbers further increased up to 40-80% after 72 hours. Possibly, cytokine-driven bystander activation (26) is involved in this further activation as we did not observe profound T-cell proliferation in the time frame of our experiments. Taken together, our model recapitulated the features of a bona fide alloreactive response.

Despite similar activation levels of CD4+ T cells and CD8+ T cells, we observed stronger induction of hydrogel contraction by CD8+ T cells. This contraction is myofibroblast-dependent, as adding only PBMCs (or CD8+ T cells) (data not shown) to the hydrogels, or selectively deleting myofibroblasts, does not induce contraction (27). Thus, the more potent contraction indicates differential immune cell-mediated activation of myofibroblasts. Recently, distinct subsets of CD8+ T cells with non-canonical functions, that divert from cytotoxic killing effect, have been described such as producing T helper cytokines or exhibiting regulatory functions (10). Canonical functions of CD8+ T cells typically involve direct cytotoxic activity against infected or malignant cells, whereas non-canonical functions include roles in immune regulation and cytokine production (28). Strikingly, no significant CD8+ T cell-mediated cytotoxicity was observed; myofibroblast viability was higher in co-culture, and 7AAD/Annexin V staining revealed minimal apoptosis or necrosis in myofibroblasts. Additionally, activation of cleaved caspase-3 and detection of double-stranded DNA breaks (γH2AX) did not indicate apoptosis (29). This suggests that cytotoxicity mechanisms are unlikely to play a role in regulating contraction, and it is more likely that the interaction between CD8+ T cells and myofibroblasts involves non-canonical functions.

Non-canonical CD8+ T cells are known to express T helper cytokines such as IL-4, IL-13, IL-6, and IFNγ. In our experiments, we found a similar percentage of CD4+ and CD8+ T cells expressing IL-4, but significantly more CD4+ T cells expressed IL-13. In contrast, a higher proportion of CD8+ T cells expressed IL-6, and overall IL-6 levels were elevated in CD8+ T cell: myofibroblast co-cultures. These findings, combined with the strong inhibitory effect of tofacitinib, suggest that IL-6 may serve as an effector molecule mediating the effects of non-canonical CD8+ T cells on myofibroblast activation and contraction. Additionally, the transfer of the contractile phenotype to naïve myofibroblasts through supernatant transfer supports the involvement of soluble mediators. This hypothesis is further supported by the observation that BFA, an inhibitor that blocks the trafficking of cytokines and growth factors from the endoplasmic reticulum to the Golgi apparatus in T cells, reduces myofibroblast activation at the gene expression level.

Mixed lymphocyte reaction (MLR) assays are essential in pre-clinical drug development for immunotherapies targeting cancer, inflammatory and transplantation-related diseases. These assays evaluate the safety and efficacy of targeting interactions between T cells and professional antigen-presenting cells (APCs) in an immunological context (30). To better understand immune-driven processes like fibrosis, it is essential to study T cell interactions with stromal cells, as these interactions are key to the orchestration of tissue adaptation. Therefore, our developed model offers added translational value compared to traditional MLR assays in studying the complex interactions between immune cells and stromal cells. These interactions are relevant in various pathological contexts, such as the pancreatic tumour microenvironment, graft-versus-host disease, and CTD pathology. SSc, a prototypic autoimmune CTD, is characterized by excessive myofibroblast activation and contraction in the skin and other affected organs. Notably, SSc is marked by a prominent influx of CD8+ T cells into the skin, especially in the early stages of the disease (31). However, it is yet poorly understood how these cells contribute to the excessive myofibroblast activation. Our work demonstrates that these cells might be directly involved in driving myofibroblast contraction. Further research into the detailed phenotype of these CD8+ T cells in SSc is needed to support this claim.

While CD4+ T cells did activate fibroblasts, their impact was less significant, likely due to differences in signaling interactions or effector functions. In many CTD-affected tissues (including SSc-affected skin), a significantly higher infiltration of CD8+ T cells rather than CD4+ T cells, particularly in early disease stages, has been observed (19, 31, 32). Cytokine-producing CD8+ T cells are likely key drivers of fibrosis, contributing to tissue destruction. Importantly, our findings are consistent with emerging evidence that CD8+ T cells can adopt cytokine-producing roles similar to CD4+ T helper subsets in certain pathological contexts (9). Possibly, this could be a consequence of myofibroblast signaling or T cell-ECM interaction, as cancer associated myofibroblasts have been shown to suppress CD8+ T cell cytotoxicity (33), which has also been suggested for tumour ECM stiffness (34).

The myofibroblast-immune cell crosstalk was directly or indirectly driven by STAT3 signaling as the JAK/STAT3 inhibitor tofacitinib strongly reduced gene and protein expression of both HLA type 2 (HLA-DR) and IL-6. In addition, TGFβ signaling, via TGFBR1, strongly affected myofibroblast activity and gene expression. Both pathways are strongly associated with fibrotic diseases including SSc (5).

Notably, a clear additive effect of inhibiting TGFβ and JAK/STAT3 signaling was observed on tissue contraction. Inhibiting these pathways had differential effects between CD4+ and CD8+ T cells. In CD8+ T cells, tofacitinib and SB-505124 had opposite effects on GZMB production, but in co-inhibition, tofacitinib was dominant in suppressing its expression. In CD4+ T cells, no such effect on GZMB expression was observed. Instead, IL-6 expression was modulated by these inhibitors in CD8+ T cells, in contrast to CD4+ T cells. These observations underscore the heterogeneity of lymphocyte responses within a signaling network. Furthermore, our findings provide mechanistic evidence on the limited efficacy of anti-IL-6 monotherapy in CTDs such as scleroderma (35, 36) and suggest that combining anti-IL6 treatment with an anti-fibrotic agent such as SB-505124 might exhibit increased efficacy in halting tissue fibrosis in CTDs. A better understanding of such differential responses will contribute to the development of more effective treatments.

While the developed 3D collagen hydrogel model provides new opportunities for translational research and high throughput drug screening, it comes along with certain limitations. Given that our model is based on MHC-driven T cell activation, we cannot exclude the possibility that the TCR-MHC interaction strength may be higher than that observed in autoimmune T cells. Since TCR signaling strength plays a critical role in determining T cell effector versus exhausted responses, our model may be biased toward the activation of certain T cell subsets. Ideally, autologous autoimmune cells should be used to validate our findings. Additionally, we did not analyse the full immune landscape, as we did not include the investigation of monocyte/macrophage phenotype and activity, despite the fact that these cell types are known to play a significant role in orchestrating fibrosis in various conditions (37). This cell type probably explains why tissue contraction was most strongly induced with PBMCs, even compared to CD8+ T cells alone.

## Conclusion

In conclusion, this study provides valuable insights into the mechanisms by which CD8+ T cells can induce myofibroblast activation and contraction, primarily through cytokine-driven activation rather than cytotoxic mechanisms. The TGFβ and JAK/STAT3 pathways were identified as critical regulators of this process, with dual inhibition demonstrating an additive effect in reducing contraction. These findings contribute to our understanding of the mechanisms underlying tissue fibrosis in systemic connective tissue diseases (CTDs). Moreover, they highlight the importance of targeting both immune and stromal cells in the treatment of fibrotic diseases.

## Acknowledgements

We thank Monique Helsen for assistance with Luminex

