## Supplemental materials and figures for "CD8+ T cells drive myofibroblast activation and contraction via JAK/STAT3 and TGFβ signaling"

### Supplemental Figures

**Supplemental figure 1**

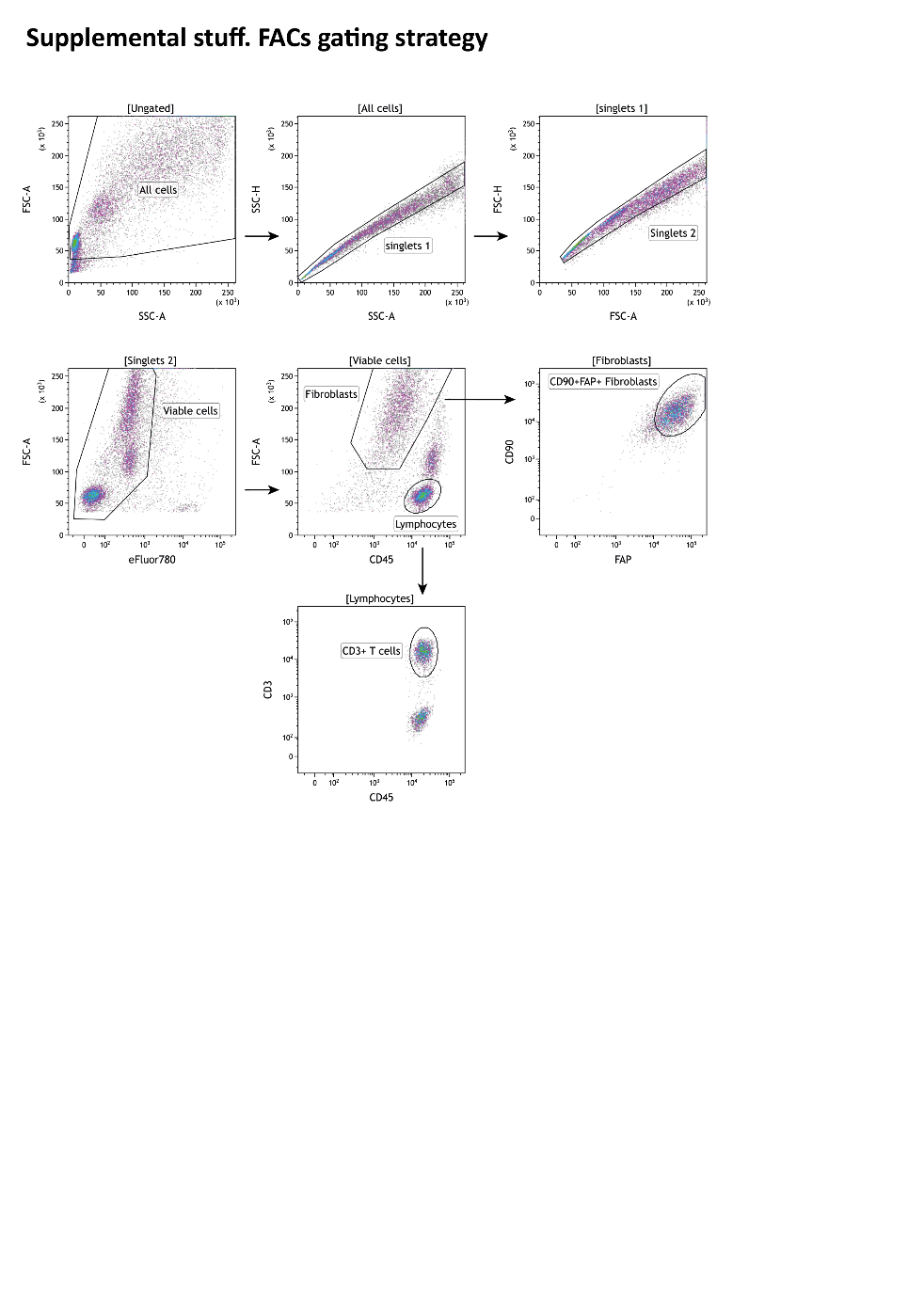

**Supplemental Figure 1. FACs gating strategy for sorting Myofibroblast and T cells for RNA analysis.** Myofibroblasts were defined as CD45^-^, and CD90^+^, FAP^+^.

FACs; Fluorescence-Activated Cell sorting.

**Supplemental figure 2**

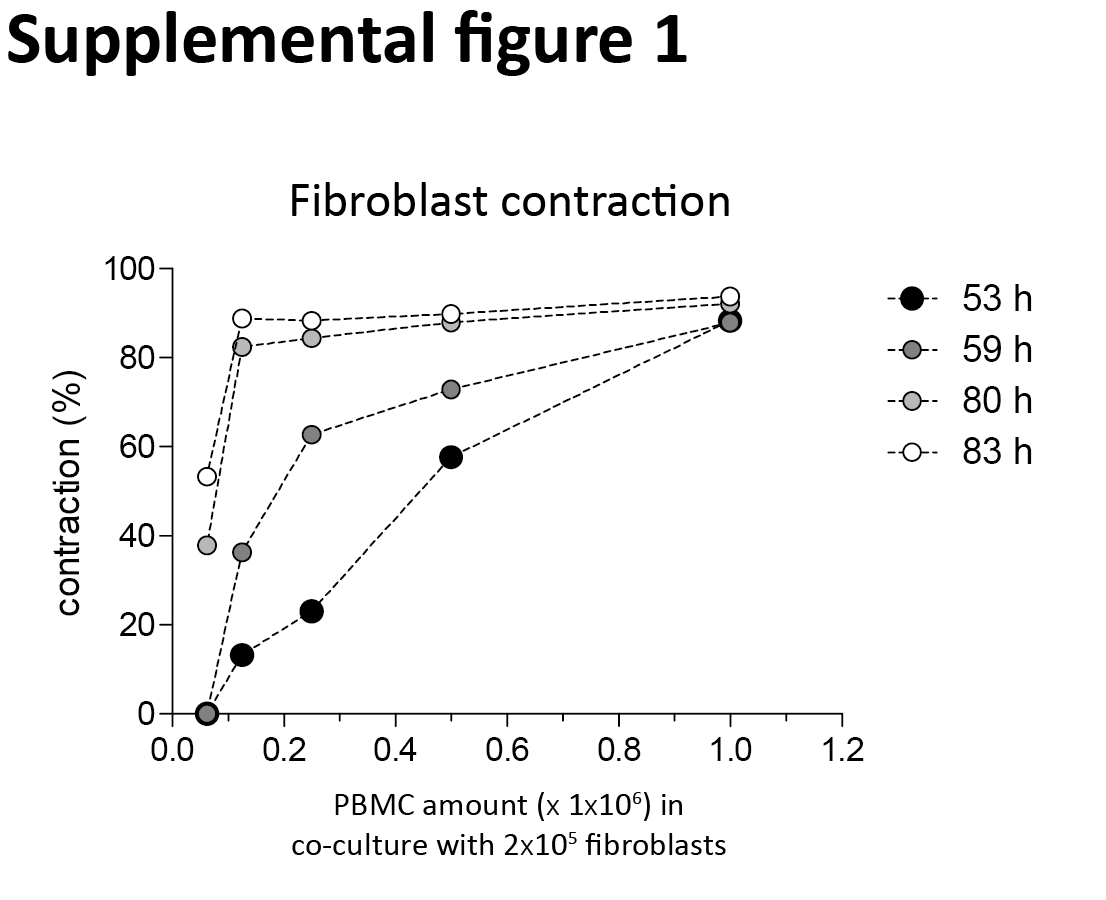

**Supplemental Figure 2. Contraction of hydrogels based on time and PBMC: myofibroblast ratio**. Primary skin myofibroblasts were co-cultured with varying amounts of PBMCs in collagen type 1 hydrogels. The extent of hydrogel contraction was measured at specific time points as indicated.

PBMCs; Peripheral Blood Mononuclear Cells

**Supplemental figure 3**

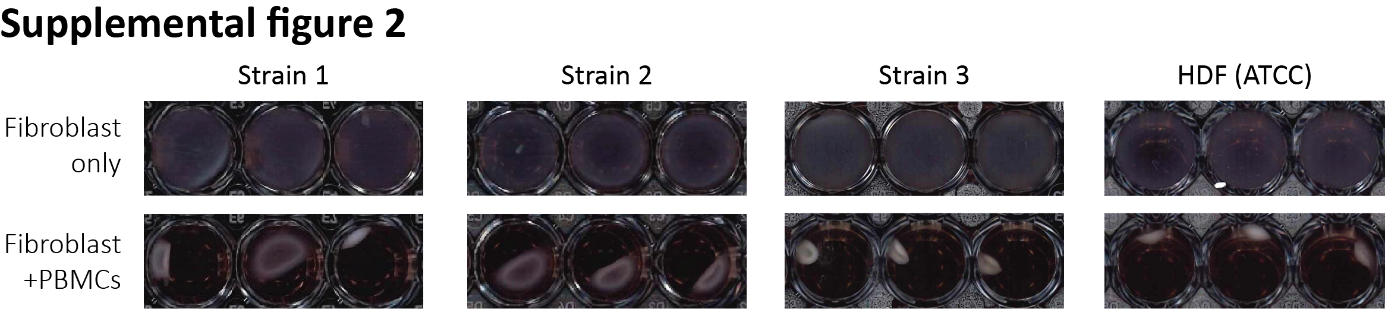

**Supplemental Figure 3. PBMC-myofibroblast co-culture induced hydrogel contraction in different myofibroblast strains.** Four strains of primary skin myofibroblasts show strong contraction after 72 h of co-cultured with PBMCs in collagen type 1 hydrogels. This includes a primary normal dermal myofibroblasts (HDF) obtained from ATCC.

PBMCs; Peripheral Blood Mononuclear Cells, HDF; Human Dermal Myofibroblasts, ATCC; American Type Culture Collection.

**Supplemental figure 4**

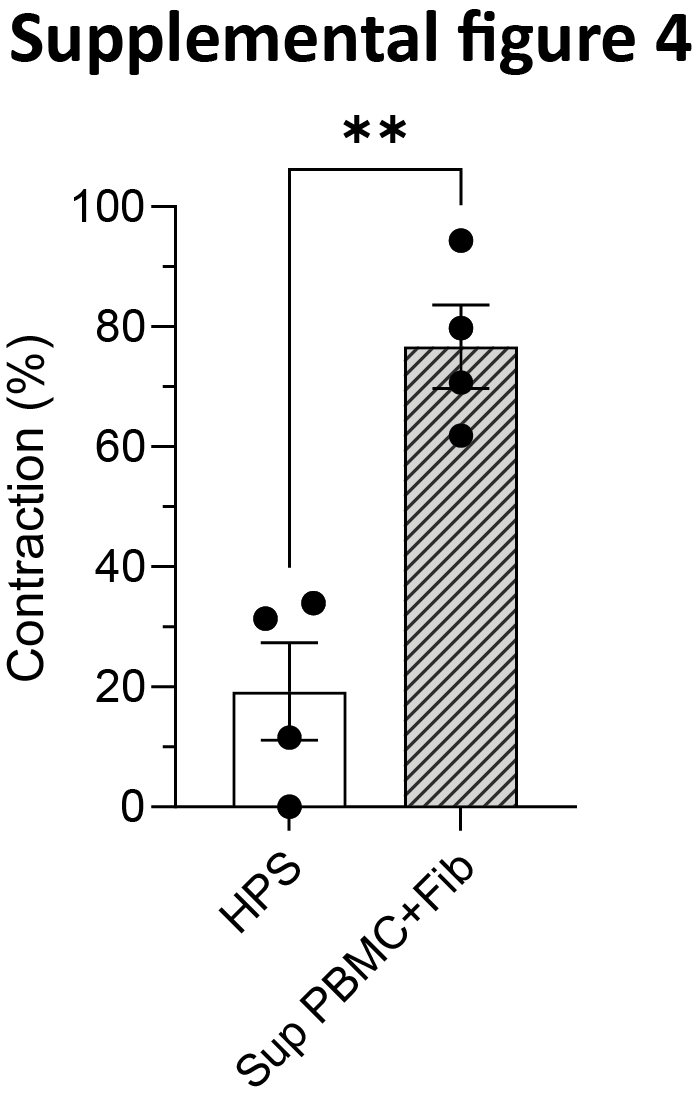

**Supplemental Figure 4. Co-culture supernatant transfer promotes hydrogel contraction in myofibroblast monoculture**. Myofibroblasts and PBMCs were co-cultured for 72 hours, after which the supernatant was collected and centrifuged twice at 300 x g. This processed supernatant (20% v/v) was then added to the medium of fresh hydrogels containing only myofibroblasts. As a control, medium that had not been exposed to co-cultures (10% HPS) was used.

PBMCs; Peripheral Blood Mononuclear Cells, HPS; Human Pooled Serum.

**Supplemental figure 5**

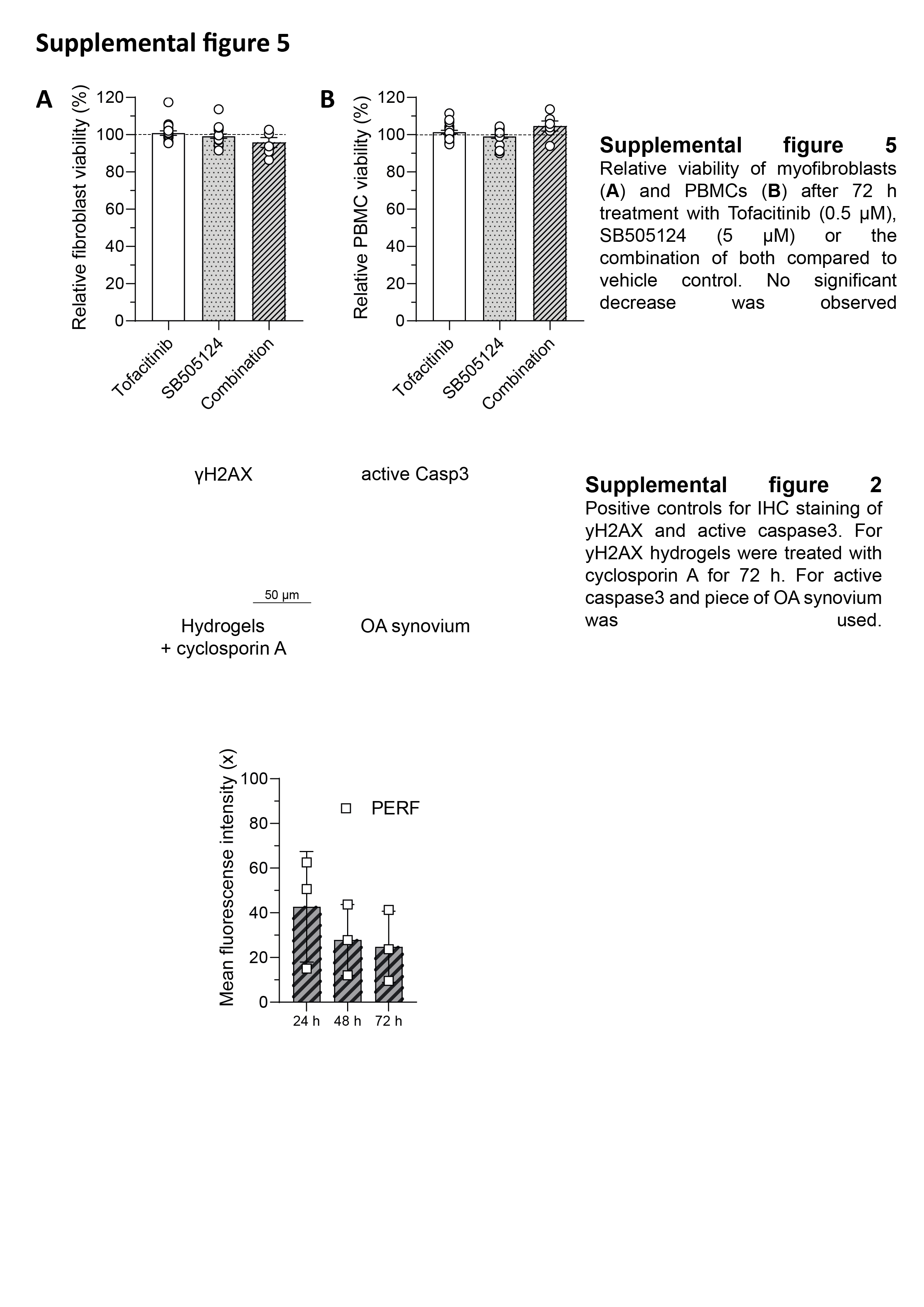

**Supplemental Figure 5. No significant decrease in myofibroblast nor PBMC viability is observed with the inhibitors’ treatment.** Relative viability of myofibroblasts (A) and PBMCs (B) after 72-hour treatment with Tofacitinib (0.5 µM), SB505124 (5 µM) or the combination of both compared to vehicle control.

### Supplementary materials

***Supplementary Table 1: List of antibodies used for immunohistochemistry***

| **Antigen** | **Clone** | **Supplier** | **Identifier** |
| --- | --- | --- | --- |
| Pro-collagen type 1 | A1/COL1A1 | Merck | ABT257 |
| alpha-smooth muscle actin | polyclonal | Abcam | ab5694 |
| Podoplanin | D2-40 | Biolegend | 916605 |
| Myofibroblast activation protein | Monoclonal | Abcam | ab207178 |
| Phospho-SMAD2 (Ser465/467) | 138D4 | CellSignaling | #3108 |
| Phospho-Stat3 (Tyr705) | D3A7 | CellSignaling | 9145S |
| Cleaved caspase-3 (Asp175) | Polyclonal | CellSignaling | 9661S |
| γH2AX | Polyclonal | CellSignaling | 9718S |

***Supplementary Table 2: List of antibodies used for cell surface flow cytometry staining***

| **Antigen** | **Clone** | **Fluorochrome** | **Supplier** | **Identifier** |
| --- | --- | --- | --- | --- |
| CD4 | RPA-T4 | PerCP/Cy5.5 | Biolegend | 300530 |
| CD3 | UCHT1 | Alexa Fluor700 | Biolegend | 300425 |
| CD8a | RPA-T8 | BV510 | Biolegend | 301048 |
| CD45 | HI30 | BV510 | Biolegend | 304036 |
| CD25 | M-A251 | PE/Cy7 | Biolegend | 356108 |
| CD134 (OX40) | ACT35 | FITC | Biolegend | 350006 |
| HLA-DR | L243 | APC | Biolegend | 307609 |
| HLA-A,B,C | W6/32 | APC | Biolegend | 311409 |
| CD90 (Thy-1) | 5E10 | APC | eBioscience | 17-0909 |
| FAP | BLR150J | APC | R&D Systems | FAB3715A-025 |
| CD69 | FN50 | Alexa Fluor700 | Biolegend | 310922 |

***Supplementary Table 3: List of antibodies used for intracellular flow cytometry staining***

| **Antigen** | **Clone** | **Fluorochrome** | **Supplier** | **Identifier** |
| --- | --- | --- | --- | --- |
| IL-2 | MQ1-17H12 | PE | eBioscience | 12-7029-71 |
| IL-4 | MP4-25D2 | PE/Dazzle594 | Biolegend | 500832 |
| IFN-γ | B27 | Alexa Fluor 700 | Biolegend | 506516 |
| IL-6 | MQ2-13A5 | Pacific Blue | Biolegend | 501114 |
| IL-13 | JES10-5A2 | PE/Cy7 | Biolegend | 501914 |
| Granzyme B | QA16A02 | APC/Fire750 | Biolegend | 372210 |
| Granzyme K | GM26E7 | FITC | Biolegend | 370508 |

***Supplementary Table 4: List of primers used for qPCR***

| **Gene** | **FWD (5’🡪 3’)** | **REV (5’🡪 3’)** |
| --- | --- | --- |
| *GAPDH* | ATCTTCTTTTGCGTCGCCAG | TTCCCCATGGTGTCTGAGC |
| *RPS27a* | TGGCTGTCCTGAAATATTATAAGGT | CCCCAGCACCACATTCATCA |
| *TBP* | GCTTCGGAGAGTTCTGGGATTG | GCAGCAAACCGCTTGGGATTA |
| *COL1A1* | AGATCGAGAACATCCGGAG | AGTACTCTCCACTCTTCCAG |
| *COL3A1* | CCTGGAATCTGTGAATCATGCC | TGCGAGTCCTCCTACTGCTA |
| *FN1* | CCCAGTCCACAGCTATTCCT | TTCATTGGTCCGGTCTTCTC |
| *FN1EDA* | TTCAGACTGCAGTAACCAACAT | GGTCACCCTGTACCTGGAAAC |
| *PLOD2* | AAGACTCCCCTACTCCGGAAA | AGCAGTGGATAATAGCCTTCCAA |
| *FAP* | GCTTTGAAAAATATCCAGCTGCC | ACCACCATACACTTGAATTAGCA |
| *ACTA2* | CTGACCCTGAAGTACCCGATA | GAGTGGTGCCAGATCTTTTCC |
| *PDPN* | GGTGCAATCATCGTTGTGGTTA | TTCAGCTCTTTAGGGCGAGTAC |
| *IL6* | AGCCCACCGGGAACGA | GGACCGAAGGCGCTTGT |
| *HLA-ABC* | TACCTGGAGAACGGGAAGGA | GTGGCCTCATGGTCAGAGA |
| *HLA-DR* | CCCTGCAGCACCACAAC | GGAACCACCTGACTTCAATGC |
| *TGFB1* | GAGGTCACCCGCGTGCTA | TGCTTGAACTTGTCATAGATTTCGTT |
